# Pooled Single-Molecule transcriptomics identifies a giant gene under balancing selection in sunflower

**DOI:** 10.1101/2021.03.17.435796

**Authors:** Hélène Badouin, Marie-Claude Boniface, Nicolas Pouilly, Anne-Laure Fuchs, Felicity Vear, Nicolas B. Langlade, Jérôme Gouzy, Stéphane Muños

**Author notes:** Corresponding author: Hélène Badouin. These authors supervized the work (Hélène Badouin, Stéphane Muños). Equal contribution (Marie-Claude Boniface, Nicolas Pouilly).

## Abstract

Genes under balancing selection control phenotypes such as immunity, color or sex, but are difficult to identify. Self-incompatibility genes are under negative frequency-dependent selection, a special case of balancing selection, with up to 30 to 50 alleles segregating per population. We developed a method based on pooled Single-Molecule transcriptomics to identify balanced polymorphisms expressed in tissues of interest. We searched for multi-allelic, non-recombining genes causing self-incompatibility in wild sunflower (*Helianthus annuus*). A diversity scan in pistil identified a gene, *Ha7650b*,that displayed balanced polymorphism and colocalized with a quantitative trait locus for self-incompatibility. Unexpectedly, *Ha7650b* displayed gigantism (400 kb), which was caused by increase in intron size as a consequence of suppressed recombination. *Ha7650b* emerged after a whole-genome duplication (29 millions years ago) followed by tandem duplications and neofunctionalisation. *Ha7650b* shows expression, genetic location, genomic neighbourhood and predicted function that provide strong evidence that it is involved in self-incompatibility. Pooled Single-Molecule transcriptomics is an affordable and powerful new method that makes it possible to identify diversity and structural outliers simultaneously. It will allow a breakthrough in the discovery of self-incompatibility genes and other expressed genes under balancing selection.

## Introduction

Understanding how and why genetic polymorphism is maintained in populations are major questions in biology. However, case studies of well-characterized loci under balancing selection are so far limited^1–7^. The genetic bases of balanced polymorphisms can be investigated with genetic mapping or molecular studies but such studies can be long and costly^8–11^. Genome scan was used as an altenative by mapping resequencing data on a reference genome in order to identify outliers^12–14^. Positive selection results in low nucleotide diversity and excess of high- and low-frequency variants, while balancing selection results in high nucleotide diversity and excess of intermediate-frequency variants. However, if alleles are too divergent, reads will not map on the reference genome, yielding false negative identification from genome scan. We hypothesized that this limit could be overcome with long-reads resequencing of transcripts, enabling alleles clustering based on their conserved domains (Figure 1). We thus conceived an experimental design consisting of pooled Iso-Seq protocol (Pacific Biosciences) sequencing. This technology produces full-lengh, high-quality sequences from individual cDNA. Because of this specificity, it should be possible to retrieve multiple alleles from a pool of individuals with a single cDNA library and to perform a whole-transcriptome scan for balancing selection in populations. Importantly, this does not require *a priori* knowledge of individual phenotypes. In addition to being affordable, transcriptomics makes it possible to focus on particular tissues and developmental stages in order to capture balancing polymporphism that are expressed in the conditions of interest. For example, in flowering plants, genetic inhibition of self-fertilization (self-incompatibility) is often controlled by a two gene-multi-allelic supergene, the genes being expressed in either pistil or the anther or pollen^2,15^.

**Figure 1.**
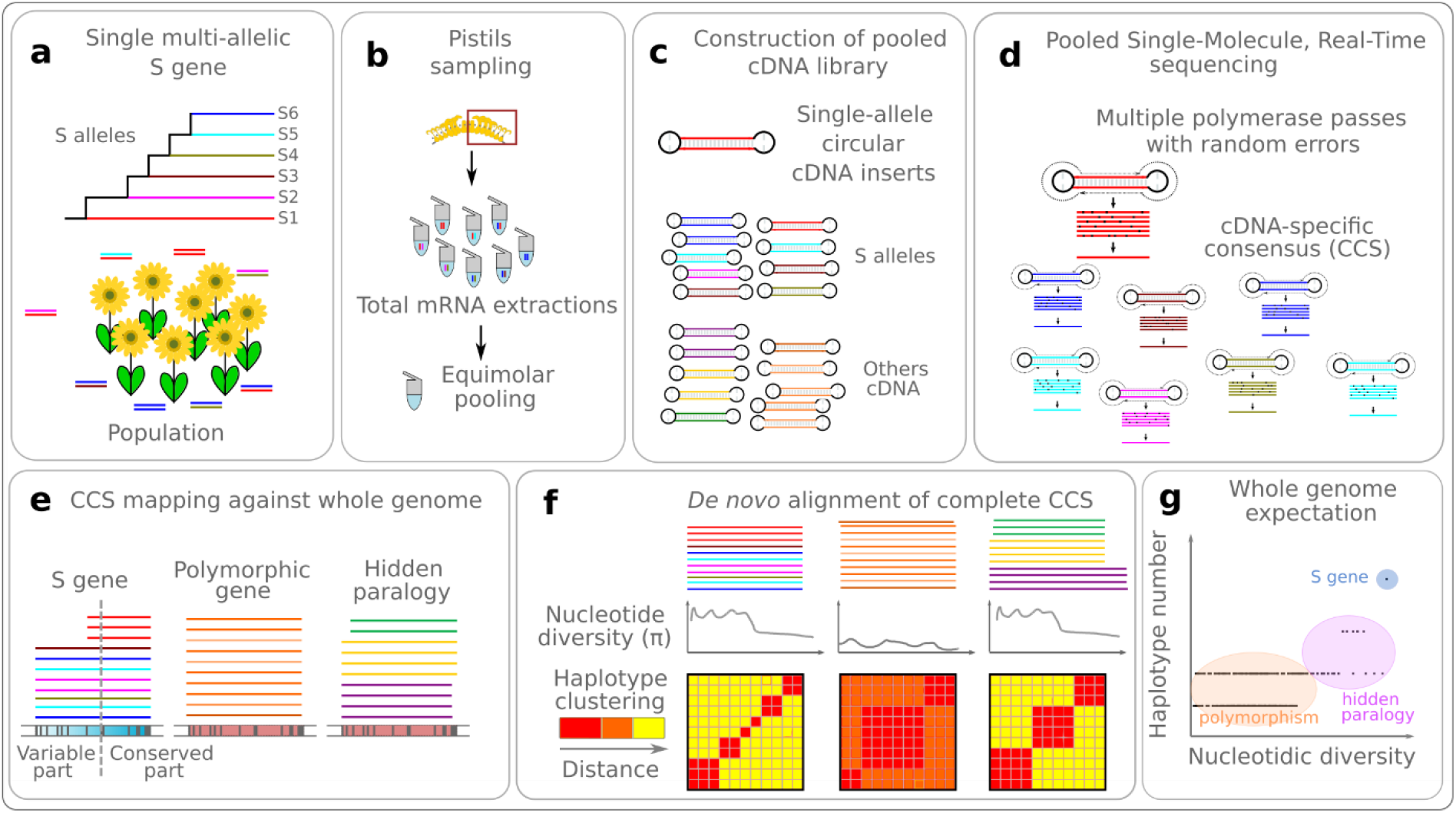
Experimental design of pooled Single-Molecule transcriptomics applied to the search of self-incompatibility genes. **a**, Example of genotypes at an S gene. As S genes evolve under negative frequency-dependent selection, even a small population can count many S alleles. **b**, Sampling of pistils and equimolar pooling of total RNA. **c**, Construction of a pooled cDNA library, with mixed isoforms and alleles of the same genes. **d**, Pooled Single-Molecular sequencing. A circular consensus sequence (CCS) is obtained for each cDNA insert, so high-quality individual allelic sequences are obtained. **e**, Mapping of CCS against a reference genome. Even highly divergent S alleles can map to the S gene locus, provided that conserved gene region can serve as anchors. If genes present in the sampled population are absent from the reference, their transcripts may map to the closest paralogue (hidden paralogy). **f**, *De novo* alignment of CCS mapping to the same genomic locus, in order to avoid underestimating diversity in transcript regions too divergent to map on the reference genome. Nucleotide diversity of an S gene is assumed to be an order of magnitude higher than genome-wide polymorphism^15^. Hidden paralogy also results in high diversity, but S genes should display more haplotypes. **g**, Expected relationships between the number of divergent alleles (haplotypes) and the nucleotidic diversity at the whole transcriptome level.

S loci, involved in self-incompatibility (SI), are excellent models for studying balancing selection because they count numerous alleles^15^. S loci form non-recombining 2-gene haplotypes that encode pistil and pollen proteins determining pollen rejection; pollination failing if pistil and pollen (or anthers) express S alleles of the same S haplotype. There is thus a selective advantage for rare alleles, resulting in maintenance of a high number of alleles over long periods of time^15^ (up to 50 alleles in a single population, Figure 1a). Furthermore, multiple independent instances of S loci exist in Angiosperms^16^. This may enable questions of evolutionary convergence in recruited genes, biological pathways and population genetics of SI to be adressed. However, S loci have been identified only in a few Angiosperm families, revealing three different molecular mechanisms^17^. Because S loci are expected to exhibit numerous alleles within single populations^15^, they are ideal models to test a long-reads transcriptomic scan. Diversity expectations for S genes have been described by Charlesworth et al. (2005)^15^. Considerable nucleotide diversity (π) should be expected, but lack of recombination leads to a limited number of highly divergent haplotypes (Figure 1f). Non-synonymous nucleotide diversity (π_n_) should be high (or higher than synonymous diversity π_s_) in domains under balancing selection.

Here we applied our Single-Molecule transcriptomics method to the search for the pistil S gene in sunflower *Helianthus annuus* (Asteraceae). The Asteraceae is the largest Angiosperm family, comprising over 2,500 species, of which 60% possess an SI system^16^. However no S genes have yet been identified despite genetic, transcriptomic and physiological studies^8,10,18,19^. Genetic control of SI in Asteraceae is similar to that in Brassicaceae^16,18^ (sporophytic SI, *i*.*e*., a gene expressed in the diploid pollen donnor controls the phenotype of pollen grains). The S locus of Brassicacae consists of two genes expressed respectively in pistil and anther and encoding a membrane receptor (SRK) and a small cystein-rich protein embedded in the pollen coat^2^. Wild sunflowers are self-incompatible, while most cultivars are self-compatible as a result of domestication and breeding^10,20^. We chose sunflower because important genetic and genomic resources were already available: a high-quality reference genome^21^, the complete repertoire of pistil- and anther-specific genes^21^ (282 and 1,530 genes respectively), and a quantitative trait locus for self-incompatibility^10^. We used our pooled Single-Molecule transcriptomic strategy on pistil from individuals of a wild *H. annuus* population (PI 413066) from New Mexico. A transcriptomic scan highlighted a giant gene under balancing selection whose expression pattern, genetic location, genomic neighbourhood and protein function are compatible with a self-incompatibility gene.

## Results

### A single balanced polymorphism expressed in pistil and colocalizing with a self-incompatibility QTL was identified thanks to Pooled Single-Molecule transcriptomics

To carry out the transcriptomic scan, we sampled pistils in 8 self-incompatible individuals in the wild population. Even in a small dataset, diversity at SI genes is expected to be high because of negative frequency-dependent selection. A cDNA library was prepared from pooled pistil mRNAs and sequenced on 4 SMRT-cells on a Sequel sequencer, yielding 1,672,707 raw polymerase reads (Supplementary Table 1). After cleaning, 707,888 consensus circular sequences (CCS) were obtained (Supplementary Table 2). In our experimental design, CCS may represent isoforms or alleles. All CCS were mapped to the reference genome, and grouped based on their intersection with predicted mRNA. We completed this by an annotation-free approach to group CCS that did not overlap significantly with predicted mRNA. The two approaches yielded 19,817 and 3,463 clusters of more than two CCS, respectively. We then aligned CCS *de novo* and measured nucleotide diversity (π) for all clusters (Figure 2a). This ensured that even highly divergent parts of the transcripts were taken into account to compute diversity. The 789 clusters with a nucleotide diversity above the 95^th^ percentile were analysed further (Figure 2b).

**Figure 2.**
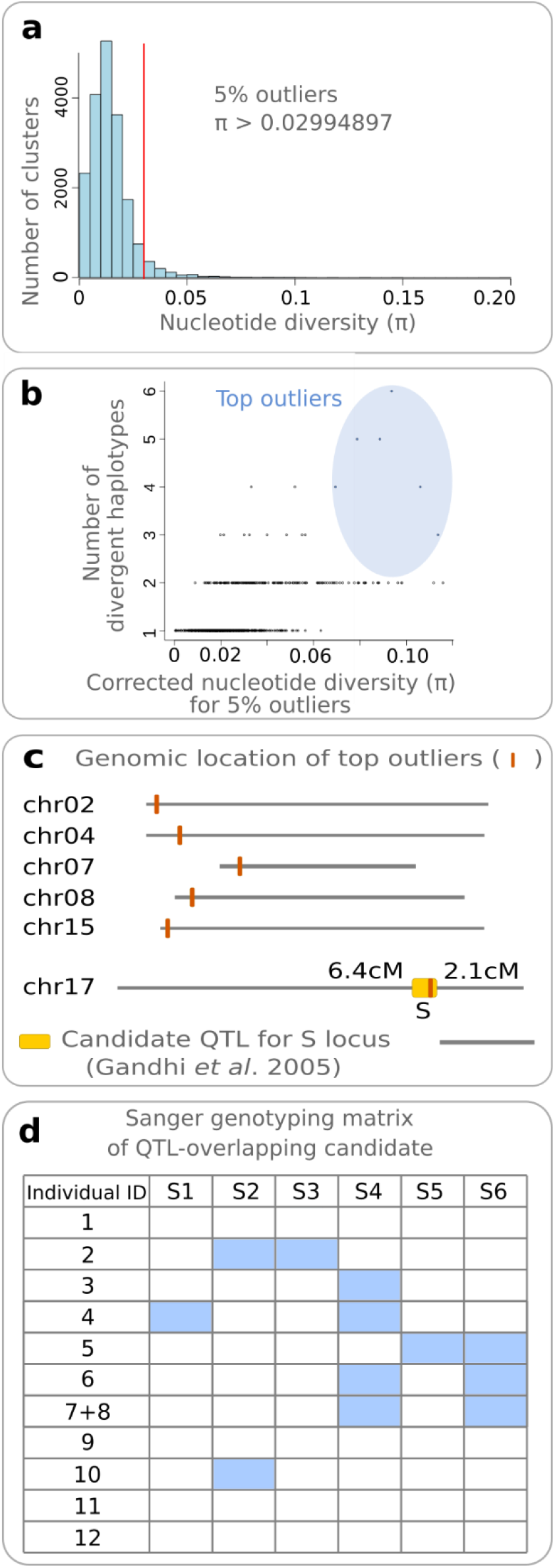
Diversity scan in pistils of wild sunflowers identifies a gene under balancing selection overlapping with a QTL for self-incompatibility. **a**, Whole-genome distribution of nucleotide diversity inferred from long-reads transcripts. The top 5% clusters were retained for further analysis. **b**, Number of divergent haplotypes versus nucleotidic diversity, computed on refined alignments (see Methods): six clusters display both a high nucleotidic diversity and three or more highly divergent alleles **c**, Genomic location of the six top outliers. A single cluster overlaps with a previously identified QTL for self-incompatibility on chromosome 17^10^. The scale bar represents 50 Mbp. **d**, Summarized genotyping matrix of the six haplotypes (S1 to S6) of the candidate gene for SI from the eight individuals used in the pooled sequencing experiments (1-8) and four additionnal individuals of the same population (9-12). Results are consistent with a one-gene model. Blue boxes indicate PCR amplification of alleles, confirmed by Sanger sequencing.

S genes are expected to display both considerable nucleotide diversity and several highly divergent haplotypes, even in a small population^15^. In Single-Molecule transcriptome data, estimations of diversity and of haplotype number may be inflated by the presence of unspliced introns or structural variants that cause misalignment. To overcome this, we used the MACSE2 software^22^ to remove non-homologous fragments, built refined alignments, computed nucleotide diversity and pairwise difference rates. We drew dendrogramms and used the same distance threshold to estimate the number of divergent haplotypes in the 789 most diverse clusters. Only 6 outliers displayed both a very large diversity and at least three highly divergent haplotypes (Figure 2b); five of them encoding proteins with putative functions (Supplementary Table 3). Among the six outliers, one colocalized with a major QTL responsible for self-incompatibility^10^, located at the top of chromosome 17 (176-191 Mb, Figure 2c, Supplementary Table 4). After manual re-alignment and cleaning, we identified 6 distinct alleles for this gene (with 2 to 7 isoforms per allele, Supplementary Table 5), displaying 2.38 to 19.51% divergence (Figure 3b).

**Figure 3.**
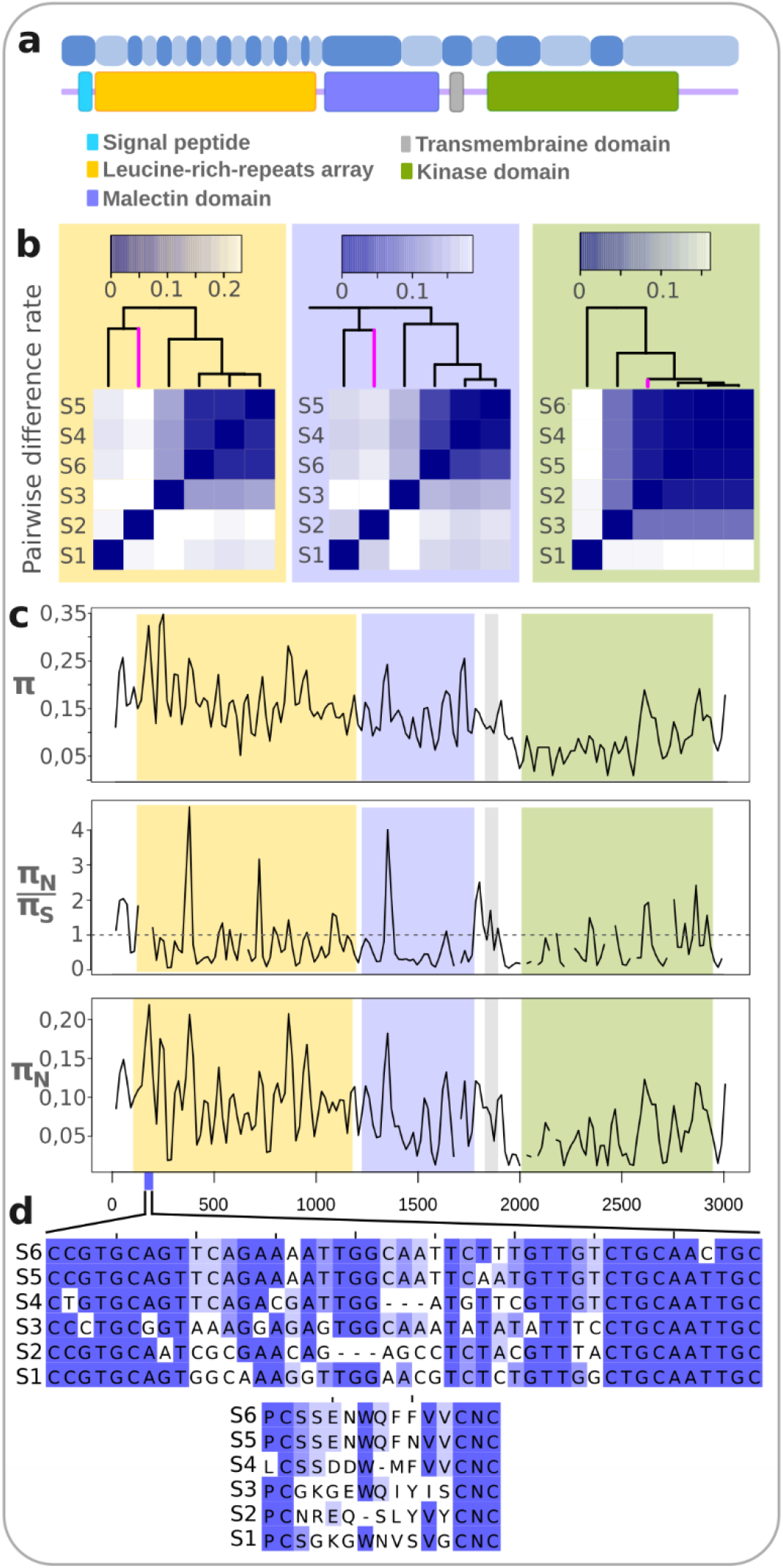
Ha7650b displays considerable inter-allelic divergence and encodes multiple hypervariable extracellular regions. **a**, Exonic structure of the *Ha7650b* mRNA and corresponding encoded protein domains. Two shades of blue are used to underline the exon limits. **b**, Matrices of allelic pairwise difference rate for mRNA regions coding for the LRR array, malectin, transmembrane and kinase domains, respectively. A recombination event between the S2 and the S4-S5-S6 branches is visible, as the S2 kinase domain (pink branch), is closer from the S4-S5-S6 group than its extracellular domain. **c**, Total (π), non-synonymous/synonymous (π_N_/π_S_) and non-synonymous (π_N_) nucleotic diversity computed by sliding windows (36 bp width, steps of 18 bp). **d**, Example of hyper-variable region in the LRR array (location shown in blue in **b**) on nucleotide (top) and amino-acid levels.

The presence of divergent alleles at one genomic locus may result from an ancient balanced polymorphism or from hidden paralogy (i.e., transcripts from distinct genes mapped at the same genomic location because the reference genome is incomplete or distant from the population studied, resulting in paralogues mistaken for alleles). This raised the risk of false positive. To verify that the alleles of the candidate gene were not hidden paralogues, we genotyped the individuals that were pooled for the transcriptomic experiment. As we studied a single population, we would expect paralogues to be amplified in all or most individuals. In contrast, in a single-gene model, each individual should carry no more than two alleles. We designed a set of specific PCR markers to amplify complete cDNA (3.2 kb) for the 6 alleles (Supplementary Figure 1a). The PCR products were sequenced with the Sanger technology to confirm the genotyping (Supplementary Figure 1b). We detected 0 to 2 alleles per individual (Figure 2d), consistent with a one gene, multi-allelic model. Furthermore, additionnal genotyping of 4 individuals of the same population detected the presence of only one allele in one of the individual (zero in the others), which suggested that we did not sampled the whole allelic diversity at this gene (Figure 2d). Hereafter, we refer to this gene as *Ha7650b* (as a reference to its orthologue in *Arabidopsis thaliana, At1g07650*, b being the paralogue identifier, see later), and to its alleles as S1 to S6. Thus, our method succeeded in identifying an expressed balanced polymorphism in the pistil from the sequencing of a single cDNA library.

### Ha7650b displays multiple hypervariable regions in its extra-cellular domains

*Ha7650b* countains 24 exons and encodes a serine/threonine kinase receptor-like, with a leucine-rich repeats (LRR) array and a malectin extracellular domain (Figure 3a). In the S locus of Brassicaceae, where the pistil gene encodes a serine receptor kinase (SRK), the residues of the extra-cellular domains that are involved in specific protein-protein recognition between the pistil and anther S genes, are under balancing selection and display hypervariability^15^. Peaks of π_N_/π_S_ or π_N_ are observed in hyper-variable domains. To test for the presence of hypervariable domains in *Ha7650b*, we measured total diversity, as well as π_N_ or π_N_/π_S._ Total diversity was higher in the transcript part coding for the extra-cellular than for the kinase domain and highest in regions coding for the LRR array. We observed at least 10 regions displaying either a high π_N_ or a π_N_/π_S_ higher than 1, that reflect balancing selection (Figure 3c), most of which were located in transcript parts coding for the LRR array, where the proportions of tri- and quadri-allelic sites reached 6.5% and 0.7% of the alignment of coding regions of the mRNA, respectively (70 and 8 out of 1,085 aligned positions respectively, Figure 3d, Supplementary Tables 6 and 7). The transcript parts encoding hyper-variable domains spanned 20-30 nucleotides of the typically 72-bp long LRR exons. This is in accordance with tri-dimentional models of LRR arrays: they form horseshoe-shaped structures, where some residues of each LRR are arranged in a concave structure that is involved in protein-protein interactions, while others play a structural role^23^. Codons encoding these residues tend be more variable than the ones encoding structural residues^24^. Thus we identified multiple small hypervariable domains probably targeted by balancing selection in *Ha7650b*. Importantly, only the part of the transcript corresponding to the malectin and kinase domain of the S1 allele mapped on the genomic locus, the LRR array being too divergent. Without using our strategy, we would have under-estimated diversity at the S locus. This highlights the power of our method to perform unbiased estimations of diversity for genes with divergent alleles.

### Ha7650b displays gigantism as a result of suppressed recombination

*Ha7650b* spans 394 kb in the reference genome and encompasses two predicted gene models (Figure 4a). All the exons of the long-reads transcripts are present in the reference genome. The size of the gene is the result of very large introns (e.g 183.8, 107.2, 52.2 or 21.7 kbp) rather than of a large number of exons. Because the reference genome was produced from a self-compatible individual^21^, this 394-kb span may have been shared with wild sunflowers or may have been a consequence of domestication or breeding. To determine the genomic structure of *Ha7650b* in wild sunflowers, we produced a draft genome assembly of a self-incompatible individual of the wild population used for the transcriptomic scan (PI 413066). Two SMRT-cells of HiFi were sequenced on a Sequel 2 (Pacific Bioscience). We obtained a single contig covering the complete *Ha7650b* locus. The *Ha7650b* transcripts mapped to a gene spanning 385 kb (S4 mapped with a coverage of 100.0% and an identity of 99.9%). The span of *Ha7650b* in PI 413066 was very close to that in the reference genome (394 kb), and very large introns were also found, although their size differed from the ones observed in the reference genome (71, 163, 82 and 47 kbp respectively, Figure 4a). Thus, *Ha7650b* gigantism predates the domestication of sunflower and the loss of self-incompatibility. In addition, the large divergence between *Ha7650b* alleles (up to 20%), as well as the differences of intron sizes between the wild and reference *Ha7650b* genomic locus, suggest that recombination is suppressed at the locus (but occasionnal recombination may occur in the kinase domain, see Figure 3a). Recombination suppressed is known to favor the accumulation of transposable elements because purifying selection is less efficient^5,25^. Therefore, our results show that *Ha7650b* is a giant gene, probably as result of long-term suppressed recombination. As annotation software use cut-off of maximum intron size to avoid spurious sequence mapping of illumina based data^26^, it would not have been possible to predict this gene model accurately without the strong evidences of single-molecule long-reads transcripts, illustrating the power of our method to reveal diversity and structural outliers simultaneously.

**Figure 4.**
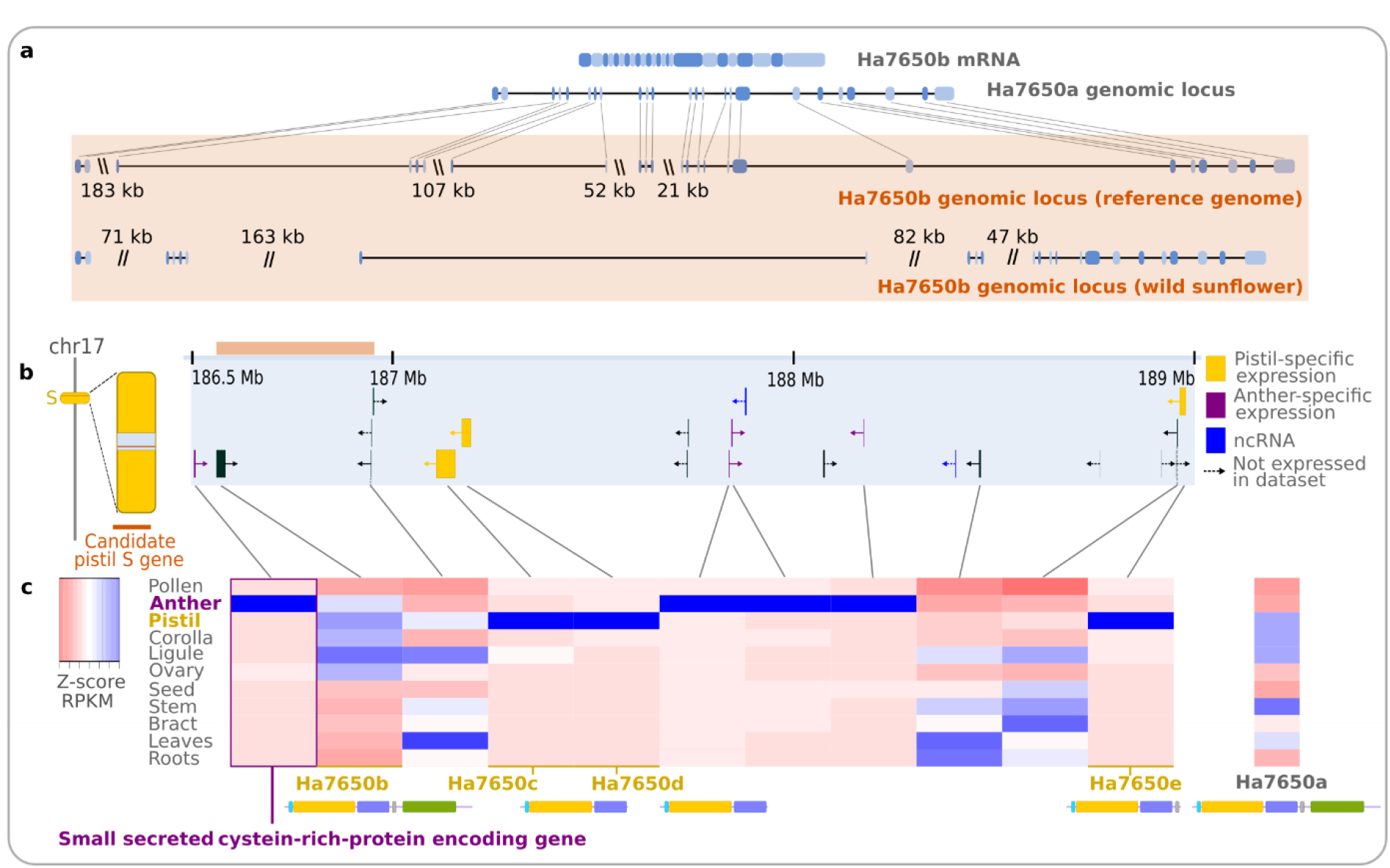
Ha7650b is a giant gene (394 kb) located in a cluster of genes specifically expressed in pistil or anther. **a**, mRNA exon structure of *Ha7650b* in a wild sunflower population and genomic structure of *Ha7650b* in the reference genome (top, 394 kb) and in a self-incompatible individual of a wild population (bottom, 385 kb). Two shades of blue are used to highlight the exon limits. Comparison with the structure of *Ha7650a* (*Ha7650b* homeologue) illustrates the size increase in *Ha7650b*. Line breaks represent long introns. **c**, Gene models predicted in a 2.5 Mb region located under the SI QTL. *Ha7650b* is wrongly split into two gene models, and several genes are specifically expressed in pistils or anthers. The bar colored in salmon indicates the limits of *Ha7650b*. **d**, Gene expression matrix in self-compatible sunflowers. Only sufficiently expressed genes are represented, the remaining ones probably being pseudogenes (see Methods). Genes specifically expressed in pistil are truncated tandem copies of *Ha7650b* (*Ha7650c*-*-e*). Among the four genes specifically expressed in anthers, one encodes a putatively small secreted cystein-rich protein and is located next to *Ha7650b*. The expression *Ha7650a* is also shown.

### Expression, genetic data, genomic neighborhood and predicted function strongly suggest that *Ha7650b* is the sunflower pistil S gene

We then sought to assess whether the balanced polymorphism that we identified may be relevant for the question of self-incompatibility. *Ha7650b* is expressed in pistil, colocalizes with a SI QTL (Figure 2c) and exhibits a large divergence between its alleles. All features are expected of an S gene^15^. To evaluate further whether *Ha7650b* meets the requirements for an S gene, we examined its genomic neighborhood and predicted function. *Ha7650b* was located within a 2.5 Mb co-expression cluster of genes specifically expressed in pistils or in anthers (186.5-189 Mb on chromosome 17, Figure 4c). In this co-expression cluster, we found 4 paralogues of *Ha7650b*, 3 of which (*Ha7650-c, -d* and *-e* respectively) were specifically expressed in the pistil, the last one probably being a pseudogene. *Ha7650c* and *Ha7650d* were tandemly located with *Ha7650b* and the two others were in its vicinity (2 Mbp), but with other genes between them. All four paralogues lacked the kinase domain, and *Ha7650c* and *Ha7650d* also lacked the transmembrane domain (Figure 4c). The four paralogues displayed a low nucleotide diversity, ruling them out as possible S genes. Among the genes specifically expressed in anther, one was adjacent to *Ha7650b* (56 kb). This gene encoded a putatively secreted small cystein-rich protein (7.26% of cystein), similarly to the anther S gene in the Brassicaceae S locus (SCR gene), located next to SRK^2^. Finally, *Ha7650b* belongs to the family of plant malectin receptor-like kinases^27^, some members of which have been shown to be involved in immunity^28^ and pollen-pistil interactions^29^ in *A. thaliana*. Thus, the genomic location, expression pattern, genomic neighborhood and predicted function of *Ha7650b* strongly suggest that it is the pistil S gene.

### Balanced polymorphism at Ha7650b emerged from ancient duplications and neo-functionalization

To understand the origin of the balanced polymorphisms at *Ha7650b*, we reconstructed its evolutive history in the Asterids (Supplementary Table 8), one of the main clade of Eudicots. *Ha7650b* belongs to the orthogroup of the *Arabidopsis thaliana At1g07650* gene (protein C0LGE0), that is present as a single copy in most Embryophytes. Our phylogenetic analysis shows that in sunflower, two duplicates (*Ha7650a* and the ancestor of *Ha7650b*, Figure 5) were retained after a whole-genome-duplication (WGD) event that occurred about 29 million years ago and is shared by most of the Heliantheae tribe (*sensus lato*, 5,000 species)^21,30^. Duplication at the *At1g07650* orthologue was shared with *Mikania micrantha (Eupatorieae)* that diverged from sunflower *(Heliantheae sensus stricto)* shortly after the whole-genome duplication. Lack of the kinase domain (the most conserved, Figure 3), makes it difficult to infer the phylogenetic position of the three functionnal truncated paralogues (*Ha7650-c*, -*d* and *e*). A maximum-likelihood tree places *Ha7650e* before the emergence of the balanced polymorphism, but with a low boostrap value. *Ha7650c* and *Ha7650d* are grouped in a branch between S1-S2 and S3-S4-S5-S6 (Figure 5). Either these two paralogues appeared independenly of *Ha7650e* from duplication of an allele of *Ha7650b* outside the non-recombining region, or the tandem arrangement of *Ha7650b*, c and d resulted in gene conversion (*i*.*e*., homologous replacement of a genomic sequence by another) between an allele of *Ha7650b* and *Ha7650c* and *Ha7650d*. The gene conversion scenario is more parcimonious, with duplication of the *Ha7650* ancestor leading to *Ha7650b* and *Ha7650e*, and duplication of *Ha7650e* resulting in a further paralogue lacking also the transmembrane domain. Internal branches between the speciation and duplication events are short, suggesting that tandem duplications and emergence of the balanced polymorphism at *Ha7650b* occurred shortly after the Eupatorieae - Heliantheae divergence. This, along with the high divergence level between the most divergent allele pairs of *Ha7650b* (19.5%), suggest that the balanced polymorphism that we identified is very ancient.

**Figure 5.**
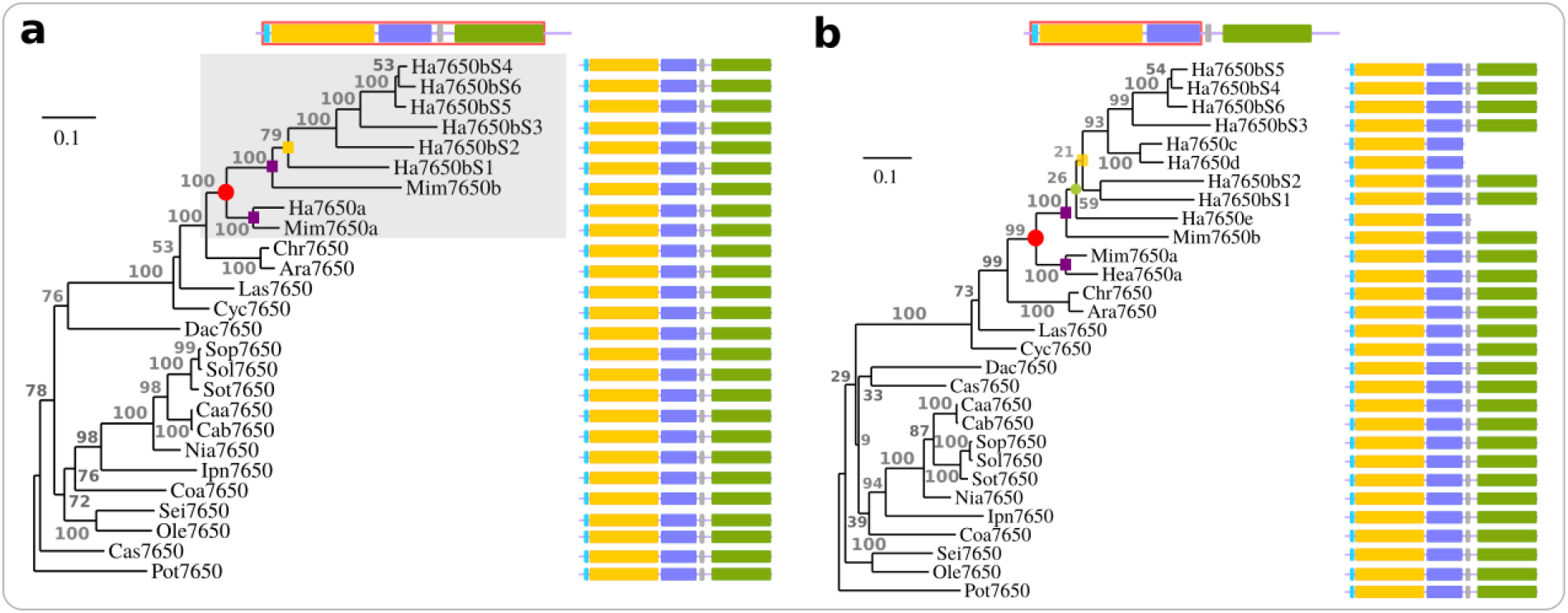
Ha7650b emerged from a complex story of duplications in sunflower. Maximum-likelyhood phylogeny of *Ha7650b* orthogroup in Asterids, with *Populus tricocarpa* (Pot7650) as an outgroup. Figures represent non-parametric branch support values. Left, phylogeny based on complete amino acid alignment and full-length *Ha7650b* orthologues. Right, phylogeny based only on the extra-cellular domains (red squared on the top gene structure), including three truncated tandem paralogues of *Ha7650b*. Red circle: 29 millions-years-old whole genome duplication (WGD). Purple square: speciation between *Helianthus annuus* and *Mikania micrantha*. Green circle: tandem duplication of *Ha7650b* ancestor.Yellow square, emergence of balanced polymorphism at *Ha7650b*. In most Asterids species, there is a single orthologue of *Ha7650b*. The *Ha7650b* ancestor gene was retained in two copies after a WGD. Several tandem duplications occurred in *H. annuus*, associated with loss of the kinase domains (green) in all duplicates but *Ha7650b*. Branch lengths and expression patterns (Figure 3) suggest neo-functionalization in the *Ha7650b-e* group after the WGD.

From a functional viewpoint, we wanted to assess which copies of *Ha7650* paralogues were most likely to have undergone neofunctionlization (*i*.*e*., gain of a new function). The function of *At1g07650* has not been investigated in *A. thaliana*, but it is expressed in several tissues (flower, fruit, leaf and root). *Ha7650a* is also expressed in several organs, including the pistil, but its greatest expression is observed in the stem (its organ-specificity index is 0.42, indicating that it is widely expressed). In contrast, *Ha7650b* is expressed in the pistil in wild *H. annuus*, and the expression of its paralogues *Ha7650c, Ha7650d* and *Ha7650e* is highly specific to the pistil in self-compatible sunflowers (Figure 4b).Moreover, the *Ha7650b-e* group and *M. micrantha* 7650b (*Mim7650b*) display longer branch lengths than the *Ha7650a* and *Mim7650a* copies in the phylogeny (Figure 5). Thus, both expression and phylogenetic patterns strongly suggest that *Ha7650a* retained the ancestral function while the ancestor of *Ha7650b-e* group underwent neofunctionalization, before the emergence of balanced polymorphism at *Ha7650b*, probably associated with the gain of a new self-incompatibility system in the Heliantheae.

## Discussion

Here we present a new method to identify expressed genes under balancing selection in populations. Focusing on pistils and sampling only eight plants of a single population, we were able to identify an ancient balanced polymorphism probably associated with suppressed recombination. The very large divergence that we observed between the four main allelic groups shows that we would not have been able to perform this analysis using the current strategies of genome scan based on short reads data. Divergence is so great in the transcript region encoding the extra-cellular domains that the short reads obtained from distant alleles would not map on the reference genome at the right genomic locus. Here the anchoring of alleles to the genome thanks to the kinase domain-encoding exons allowed their identification. Furthermore, *de novo* alignment of alleles made it possible to obtain an unbiased estimation of diversity in the exons encoding the LRR array, while reliance on short-read mapping would yield missing data. Surprisingly, the gene that we identified spans almost 400kb. Detection of such a large gene, split into two genes in the annotation of the reference genome, with several unannotated exons, would have been very difficult or almost impossible without long-reads transcript evidences enabling the delineation of the true gene boudaries. Our method thus allows extracting simultaneously interesting diversity and structural gene features.

Beyond the novelty of our method, the balanced polymorphism that we identified is likely important on an evolutionary and functional viewpoint. Our results strongly suggest that we have found the pistil self-incompatibility in sunflower. The ancient origin of the balanced polymorphism is nevertheless suggestive of an important function. The *At1g07650* orthogroup is single-copy in most Embryophytes, but was retained as duplicates and underwent neofunctionalization in one copy after whole genome duplication in the Heliantheae tribe (5,000 species). A more complex story of tandem duplication precluded the emergence of balanced polymorphism in the Heliantheae (*sensus stricto*), shortly after its divergence with *M. micrantha*, hence the balanced polymorphism may be shared by hundreds or thousands of species.

Our pooled long-reads-transcriptomics scan was very powerful to scan for highly polymorphic genes in a population. It is particularly powerful for genes that possess numerous^15^ and/or highly divergent alleles^31,32^, and should allow considerable acceleration in the discovery of expressed genes under balancing selection. By targetting particular tissues or developmental stages, it makes it possible to focus on genes expressed in the conditions of interest. Since it depends on sequencing only a single cDNA library, it is affordable and easy to carry out. Furthermore, it is not necessary to know the phenotype of the individuals that constitute the pool. This is very useful for the study of phenotypes that are difficult to characterize, such as self-incompatibility^8^, where distinct incompatibility profiles are not associated with morphological differences in flowers. Because of this versatility, pooled Single-Molecule transcriptomics will have multiple fields of application, and will allow rapid discovery of new genes under balancing selection, such as immunity genes, sex chromosomes (X/Y or Z/W) or self-incompatibility loci.

## Online methods

### Data availability

Single-Molecule sequencing data (IsoSeq and genomic) have been deposised to the NCBI Sequence Read Archive (PRJNA603280 and PRJNA603997 respectively). The Locus *Ha7650b* of the wild sunflower PI413066 was submitted to NCBI Genbank under accession MN990444.

### Graphs and statistics section

Unless stated otherwise, statistical analyses and graphs were carried out with R version 3.4.4^33^.

### Generation of pooled Single-Molecule, Real-Time transcriptomics datasets

#### Plant material

In 2017, four populations of wild *H. annuus* maintained in the sunflower gene bank (CRB) at the Laboratory of Plants-Microorganisms Interactions (LIPM), INRAE, Toulouse, were grown in an insect-free greenhouse (8 to 12 individuals per population). Self-incompatibility was phenotyped qualitatively by checking the absence of viable seeds in inflorescences. In one of the populations for which no seeds were observed (PI 413066 from New Mexico), pistils from diverse developmental stage (from central immature disc florets to well-opened peripheral disc florets) were dissected, frozen in liquid nitrogen and stored at - 80°C.

#### mRNA extractions and Single-Molecule, Real-Time sequencing

Total RNA were extracted using a QIAZOL lysis reagent (QIAGEN, USA) following manufacturer’s protocol. Total RNA was further purified by an additional purification with the NucleoSpin® RNA Clean-up kit (Macherey-Nagel, Germany). Total RNA was treated with TURBO DNA-free kit (Thermo Fisher Scientific, USA) for 30 min at 37°C to remove residual DNA. RNA concentration and purity were checked with a NanoDrop Lite Spectrophotometer (Thermo Fisher Scientific, USA). RNA integrity was assessed by the Agilent 2100 bioanalyzer, using the Plant RNA Nano chip assay (Agilent Technologies, USA). Six individual samples and a pool of two individuals (for which less raw material had been available) were pooled in an equimolar way. The GTF platform of the University of Lausanne prepared cDNA libraries and sequenced them on a Sequel SMRT sequencer. One library was prepared with a standard magbead protocol (with no size selection) and a second with a modified protocol that optimizes yield. Each library was sequenced on 2 SMRT cells with a 600 minutes reading time (batch 1 and batch 2 respectively, Supplementary Table 2). A second library was prepared from the same pool of total mRNA, using a modified protocol that optimizes yield, and also sequenced on 2 SMRT cells.

### Scan of diversity using Single-Molecule, Real-Time transcriptomics

#### Obtention and mapping of consensus circular sequences (CCS)

We applied the css (default parameters) and lima (--isoseq, primer_5p: AAGCAGTGGTATCAACGCAGAGTACATGGGG; primer_3p:AAGCAGTGGTATCAACGCAGAGTAC) software of SMRT Link 6.0.0 (https://www.pacb.com/support/software-downloads/) to extract, and clean up circular subsequences (CCS). We obtained 188,154 and 519,734 high-quality CCS for batches 1 and 2 respectively. The 707,888 high-quality CCS were mapped against the sunflower XRQ reference genome v1 (www.heliagene.org, Genbank Accession MNCJ00000000) with gmap^34^ (version 2018-07-04, indexing: gmap_build -q 2; mapping: --allow-close-indels=1 --max-intronlength-middle=500000 --cross-species. Hits were found for all but 6,425 CCS. CCS that mapped at multiple locations (5.61%, *i*.*e*. 41,819 CCS) were removed. In total, 659,644 high-quality CCS were uniquely mapped.

### Clustering of CCS

Alleles divergent from reference sequences may map only partially. In order to perform unbiased measures of nucleotidic diversity, it was necessary to align CSS *de novo*. We first defined groups (or clusters) of CCS that mapped to the same genomic locations. Long-reads transcripts include up to 15% of pre-mRNA with remaining introns, transcripts with long UTR that overlap with adjacent genes or read-through (i.e., transcripts resulting from transcription of two adjacent genes). Because of this structural complexity, a particular care had to be taken to avoid clustering non-homologous sequences, which would result in a false signal of high diversity.

#### Clustering based on overlap with predicted exons

We first grouped CCS that overlaped with predicted mRNA. We used *bedtools*^35^ *intersect* to count the number of overlapping nucleotides between each exon of predicted mRNAs (Eugene annotation v1.1) and exons inferred by gmap for the CCS. Only stranded overlaps (*i*.*e*., on the same strand) were considered. Exon overlap values were propagated at the mRNA level to compute reciprocal overlap rates between predicted mRNAs and CCS. We associated CCS with a predicted mRNA if at least 60% of its length overlaped with the predicted mRNA, or conversely. We chose this threshold to account for possible prediction mistakes in the genome annotation, and because a fraction of CCS represent shorter transcript variants. Similarly, a CCS could be associated with several mRNAs, to account for structural differences between predicted mRNA and CCS (e.g., CCS spanning two or more adjacent gene models). For each predicted mRNA, a group of CCS was formed. Groups that were totally included in another were removed. This yielded 24,075 clusters of CCS, including 19,817 with more than two CCS.

#### Clustering of CCS not overlapping with predicted exons

In the first step, we left aside 45,728 CCS that were uniquely mapped on the genome but did not sufficiently overlapped with a predicted mRNA. As some genes may be unannotated or misannotated, we performed an annotation-free clustering. First, we used *bedtools cluster* to perform a preliminary clustering (grouping any overlapping CCS in a stranded way without considering overlap rate). Within each group, we computed pairwise reciprocal rates of overlap. Then, we associated two CCS if the first pair member had a minimum rate of overlap of 0.6 with the second, or conversely. We built a graph of associations and extracted subgraphs where every pair of CCS were associated. Redundancy was removed by suppressing clusters fully included in others. This second clustering step yielded 3,463 clusters with more than two CCS. In total, we obtained 23,280 clusters with more than two CCS.

### Diversity scan

In our experimental setting, a single gene can be represented by several CCS that represent either isoforms, transcriptional variants, alleles, or paralogues in cases of hidden paralogy (i.e., if a transcript procuded by a gene wrongly map to another gene). Hereafter we refer to CCS that may be alleles or paralogues as haplotypes. In order to remove introns and long UTR that may cause alignment errors, non-homologous fragments were removed with the sub-program trimNonHomologousFragments of MACSE v2.03^22^ (-min_cov 2 -min_trim_ext 20 -min_trim_in 40). After trimming, 18,696 CCS clusters with at least 3 CCS remained. CCS clusters were aligned with mafft v7.307^36^ (default parameters). Nucleotide diversity (π) was computed with the eggstats^37^ (parameters groups=no outgroup=no multiple=yes minimum=0.2 coding=no). Clusters with a nucleotide diversity (π) above the 95^th^ percentile (769 CCS clusters with π >=0.299) were further processed.

#### Refined alignment and error correction

Alignement errors, presence of low-quality CCS, remaining introns or untranscribed transcript regions (UTR), can inflate estimations of diversity and haplotype number. Thus clusters were re-aligned with mafft, using recommanded parameters for sequences with structural variations, and reduced penalties for gap opening (-- ep 0 --genafpair --maxiterate 1000 --op 0.5 --lop -0.5). Low-quality CCS display high rates in insertions and deletions (INDEL). Single-nucleotide gaps caused by a single CCS in multiple alignments are most likely artefacts of Single-Molecule, Real-Time sequencing. We computed individual rates of 1-bp insertions or deletions (INDEL) from multiple alignments. Visual examination of several multiple alignments helped defining a threshold above which poor-quality CCS caused alignment issues (e.g., close insertions and deletions compensing each others, not detected by alignment software and resulting in erronous single nucleotide polymorphisms). CCS with an INDEL rate higher than the 90^th^ percentile (0.010468042) were removed. In addition, we performed two rounds of INDEL corrections: insertions affecting single CCS were removed whatever their size. Individual 1-bp deletions were corrected: for monomorphic sites (*i*.*e*., a unique nucleotide at that position), gaps were replaced by the nucleotide, at polymorphic sites they were replaced by ‘N’ to restore phase. Nucleotidic diversity was re-computed as previously.

#### Estimation of the number of divergent haplotypes

We computed pairwise difference rates (*i*.*e*., the number of difference divided by number of aligned bases) between CCS. Matrices of difference rates were converted to distance matrices without transformation, and a hierarchical clustering was performed using the hclust function (complete linkage method). A common threshold of 0.1 was applied to cut dendrograms (i.e., trees resulting from the hierarchical clustering) in all clusters with the cutree function. To estimate the number of divergent haplotypes, we counted the number of groups in the tree at the cut level. We plotted the number of haplotypes as a function of nucleotidic diversity (π). A group of six outliers (nucleotidic diversity >= 0.07 and at least three haplotypes) was analysed further.

### Analysis of the 6 top diversity outliers

#### Manual curation of the alignments

We removed remaining non-homologous regions (UTRs and a few remaining introns) that may inflate the estimation of diversity and haplotype number, as well as remaining artefactual INDEL. Based on a preliminary visual clustering, we were able to correct 1-bp deletions inside each haplotypic group (*i*.*e*., if a position was monomorphic inside an haplotypic group, the gap was replaced by the corresponding nucleotide rather than by an ‘N’, even if it was polymorphic at the level of the whole alignment).

#### Non-synonymous and synonymous diversity

We re-aligned outlier clusters in respect with phase with MACSE, and extracted coding regions with NCBI blastp and ORFinder (https://www.ncbi.nlm.nih.gov/orffinder/). When CCS displayed different limits for coding regions, only regions that were coding and in the same frame for all CCS were considered. At that stage, one cluster was excluded because it consisted in non-coding sequences that mostly mapped in an intronic region. Coding and non-coding diversity were computed on complete alignements and by sliding windows (18-nucleotides width, 9-nucleotides step).

#### Overlap with QTL for self-incompatibility

A major self-incompatibility QTL had been identified in Gandhi et al. (2005)^10^. The two closest Sequenced Tag Site markers (ORS735 and HT925) were retrieved respectively from NCBI (accession number BV006734.1) and from the sunflower transcriptome database (https://www.sunflowergenome.org, Ha412T4l22256C0S1). Both sequences were mapped with the blastn programme of the blast+ suite^38^ (default parameters) against the sunflower genome. Coordinates of the best hits were used to define the physical limits of the QTL. A single outlier was located within this interval.

### Genotyping and Sanger sequencing of Ha7650b

The number of *Ha7650b* alleles was estimated to six (cf. below, fine-scale diversity analysis of *Ha7650b*). We genotyped each individual used in the sequencing pool, as well as four additionnal individuals of the same population, for which pistils had been sampled in the same conditions. Specific PCR primers were designed to amplify full-length cDNA from 5’ to 3’ UTR for each putative S allele. Three pairs of primers per allele were also designed to amplify overlapping 1-kb amplicons, such as the combinations of 1-kb amplicons was specific of each allele. The 1-kb amplicons were generated from purified PCR product of full-length amplification of cDNA. PCR conditions are detailed in Supplementary Methods. They were then sequenced with the Sanger technology. Sanger sequences were aligned with *Ha7650b* alleles to validate long-reads sequences and identify the amplified allele.

#### Detection of genes specifically expressed in pistils or anthers

We used index of organ-specificity computed in Badouin et al (2017)^21^ from RNAseq data in 11 vegetative and floral organs. An index of expression specificity (τ^39^) had been computed for each gene. Organ-specific genes were defined by a τ higher than 0.8, the organ in which the expression was highest was retrieved.

### Fine-scale diversity analysis of Ha7650b

#### Assessment of completeness of the CCS dataset

To identify potential false negative (i.e., *Ha7650b* CCS that may not have mapped to Ha7650b genomic locus), we blasted with blastn or blastp nucleotide or inferred protein sequences of two CCS (S6i1 and S4i1, Supplementary Table 5) against the database of all CCS. We identified three additional CCS that were highly similar to other two CCS of *Ha7650b* (S1i1 and S1i2). This yielded a final dataset of 25 CCS (including two CCS that displayed high INDEL rates, Supplementary Table 5).

#### Manual curation of the alignment

We performed again manual curation and INDEL correction, as described above for the study of the top diversity outliers. Despite this curation, translation of CCS showed that frameshifts remained in several sequences. Thus the sequences were re-aligned with MACSE alignSequences (reducing to 10 the cost for a stop codon), and finally the MACSE alignment was manually corrected, obtaining a final alignment size of 3,026 bp (after removal of non-homologous 5’ and 3’ UTR).

#### Determination of the limits of exons and of functional domains

In order to determine the limits of exons, we used the exons inferred by mapping of the CCS by gmap against the reference genome, as well as the predicted exons in the reference genome (Eugene and ncbi RefSeq annotations). Comparaison with paralogues confirmed these limits (malectins are known to be structurally conserved^27^). In order to infer the limits of functional domains, corrected CCS sequences were translated, and translations used as input for blastp or InterProScan^40^. The largest estimation was retained for the limits of the different domains (signal peptide, LRR array, malectin, transmembrane and kinase domains). Subsequent analyses were performed on the complete alignment of sub-alignments corresponding to CDS regions encoding the respective functional domains.

#### Estimation of the number of alleles

Matrices of pairwise differences were computed with a custom python script and displayed as heatmaps. We inferred the presence of six putative alleles in the dataset, hereafter referred to as S1 to S6. Subsequent analyses were performed either on the alignment of all 25 Ha7650b CCS, or on an alignment of one representative CCS per allele. A representative CCS of each allele was retained for further analysis for S1, S4, S5 and S6. For S2 and S3 respectively, two CCS displaying complementary splicing variations were used to build a composite sequence covering the whole transcript.

#### Measure of total, synonymous, non-synonymous diversity

The total, synonymous and non-synonymous diversity in the alignment were measured with egglib v3^37^. This was done on the complete alignment and by sliding windows (width 36 bp, step 18 bp). The number of bi-allelic, tri-allelic and quadri-allelic position was computed with a custom python script.

### Sequencing and assembly of the Ha7650b locus of the PI413066 wild sunflower from New Mexico

DNA was extracted from youngest leaves from 6-weeks old plants grown in an insect-free greenhouse using the Mayjonade et al. protocol^41^. At the end of flowering, the number of viable seeds per square cm was measured in all inflorescences to ensure the selection of a self-incompatible plant, and the plant with the lower mean seed number was chosen for sequencing. Two HiFi libraries were prepared and sequenced on Sequel 2 by the gentyane platform according the Pacific Biosciences specifications. The ccs software of SMRT Link v8.0.0 (-- min-passes 4 --min-length 5000) was used to extract 2,367,365 circular consensus sequences (N50: 14.2kb) corresponding to a total of 32,937,296,594 nucleotides. Two assemblies were first performed with CANU^42^ version 1.9 or FALCON^43^, with a filtering step of falcon replaced by til-r software (N50: 533,277 and 1,348,425 bp respectively). Then, to assemble the *Ha7650b* locus, was performed with Minimus2 (default parameters, version amos-3.1.0). Minimus2 found an overlap of 60,821 nucleotides at 99.82 percent identity and generated a consensus sequence spanning the whole locus in a single contiguous sequence. The mRNA sequence data were mapped using gmap (version 2019-09-12, same parameters as above) to annotate a gene spanning 385,712 nucleotides in the PI413066 wild sunflower. Parameters are provided in Supplementary Methods.

### Evolutionary history of *Ha7650b*

#### Search for paralogues of *Ha7650b* in the sunflower genome

##### blastn and blastp were used to search for paralogues of Ha7650b

Three truncated tandem duplicates of the *Ha7650b* were also detected with blastn in the physical proximity of *Ha7650b*, and lacked the gene part encoding the kinase domain. Moreover, the closest full-length predicted paralogue to *Ha7650b* was located on chromosome 1 in the version 1 of the assembly (HanXRQChr01g0028671).

### Phylogeny of the *Ha7650b* in Asterids

#### Dataset building

The homeologue of *Ha7650b* was annotated^21^ as the orthologue of the *At1g07650* gene. This was confirmed with blastp of the amino acid-translation of *Ha7650b* alleles. The amino-acid and CDS sequences of members of the *At1g07650* orthogroup in Asterids were retrieved from OrthoDB^44^, and *Populus trichocarpa* was used as an outgroup. In addition, we performed blastp against the predicted proteomes of recently published Asterids genome (*Artemisia annua*^45^, *Chrysanthemum nankingense*^46^, *Panax notoginseng*^47^, *Calotropis gigantea*^48^, *Gelsenium sempervirens*^49^ *and Mikania micrantha*) in order to retrieve additionnal orthologues. Since this orthogroup mostly counts single-copy genes, and malectins receptors have been found to be highly conserved on a structural point of view^27^, we retained orthologues only when they were full copy, in order to avoid annotation issues. The exception was the genome of *Mikania micrantha*, a member of the Heliantheae tribe that shares a whole-genome duplication with sunflower. We performed a manual annotation by tblastn of the predicted amino-acid sequences of S1-S6 and Augustus-online^50^. Two full-length homeologues were retrieved in the two syntenic regions resulting from the whole-genome duplication. The final dataset included 28 Asterid and 1 outgroup sequences (Supplementary Table 8).

#### Codon and amino-acid alignment

We aligned the predicted CDS sequences with mafft v7.307^36^ (default parameters). We extracted sub-alignments corresponding to the main functional domains (LRR array, malectin, transmembrane and kinase domains), and for the whole extra-cellular domain. This was done with or without the sunflower tandem repeats that lack the kinase domain. As programs that filter alignments may mistake hypervariable domains for poorly aligned regions, the alignments were filtered manually. This was facilitated by the fact that the *At1g07650* orthogroup displays a strong structural conservation and that hypervariable domains were surrounded by more conserved regions. The more divergent 5’ and 3’ parts of CDS alignments had to be trimed and clade-specific insertions were removed, as not informative. CDS alignments were translated to obtain amino-acid alignments. Direct use of amino-acid sequences gave similar alignments.

#### Tree reconstruction

The best models of nucleotide or amino-acid substitution for each alignment were determined with ModelTest-NG^51^, and were the Jones-Taylor-Thornton (JTT) for all amino-acid alignments and GTR+I+G4 for codon alignments. PhyML^52^ was used to reconstruct the trees and TreeDyn^53^ to vizualise them. To compute branch support values, 100 non-parametric boostraps were carried out.

## Legends of Supplementary Figures

**Supplementary Figure 1:**
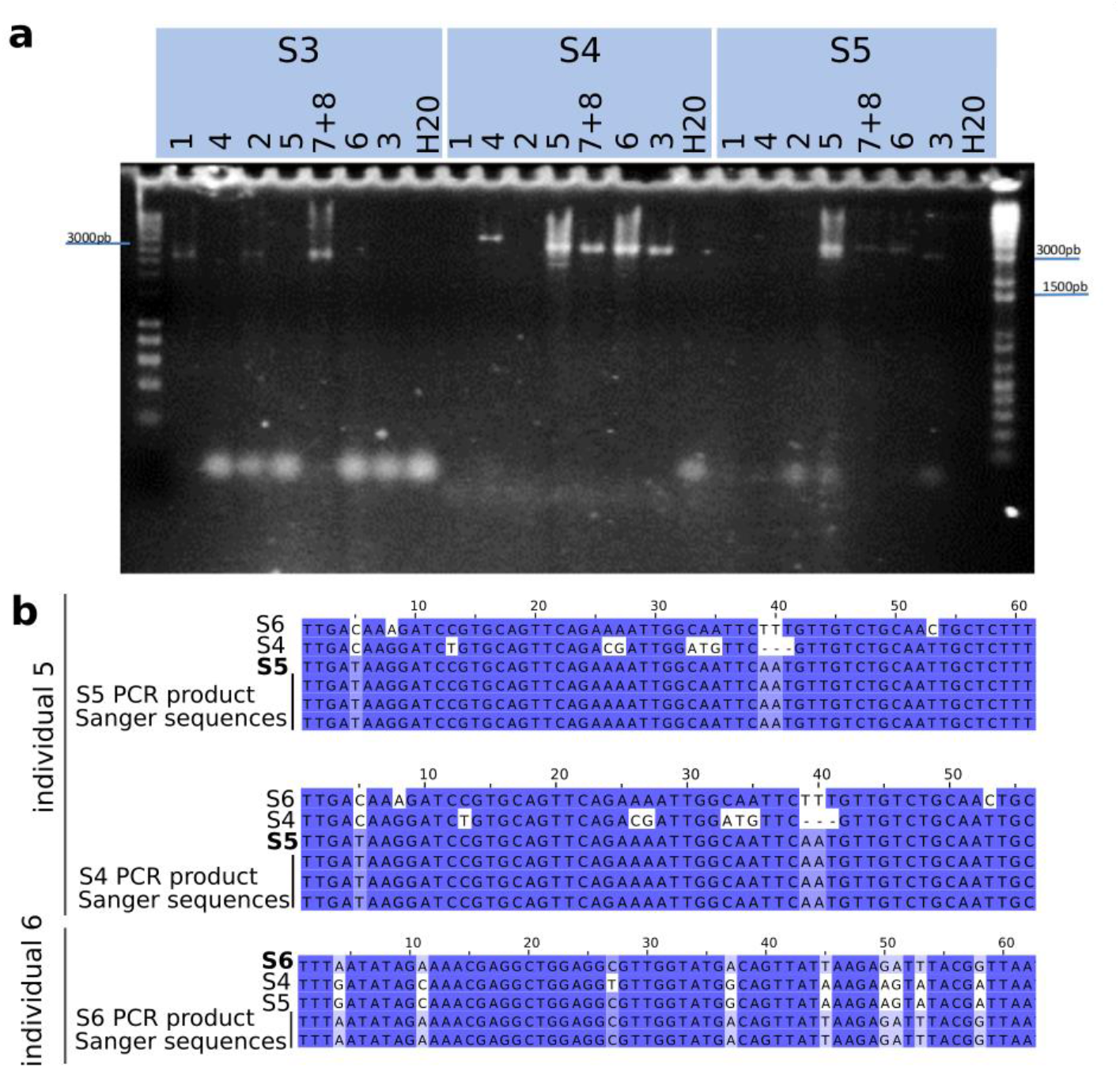
Long-range PCR and Sanger genotyping of S alleles. **a**, Long-range polymerase chain reactions (PCR) of S3, S4 and S5 alleles from total cDNA of each of the sample included in the transcriptomics pool. 1-6 are single individuals and 7+8 refers to a pool of two individuals. Specific primer pairs were designed for each allele. Individuals 1, 2, 3, 4 and 6 appear to have no more than two putative alleles (respectively no more than 4 in the 7+8 pool), consistently with a one gene model. However, in individual 5 amplification is observed for S4, S5 and S6 (not shown), but this results from non-specific amplification (**b**). **b**, Comparison of CCS and Sanger genotyping of S4, S5 and S6 PCR products, with two examples of true positive genotyping (top and bottom pannels) and one example of false positive (middle pannel). In individual 5, S4 primers actually amplified S5 sequence. Therefore individual 5 carries only S5 and S6 sequences, consistently with a one gene model.

## Acknowledgements

This work was supported by starting grants from Université Claude Bernard Lyon1 (BQR) and Université de Lyon (IDEX/IMP/2017/17 and IDEX/ELAN/2018/07) to HB. We thank the Lausanne GTF and the INRAE Gentyane platforms for producing the Single-Molecule transcriptome and genome data, respectively. We thank the GenoToul bioinformatics platform for computing resources. We thank Patrick Vincourt for discussions, advices and for feedback on the manuscript.

## Authors contributions

HB conceived the research. MCB, FV, NL and HB chose plant material. MCB and HB obtained plant material and NP obtained high-quality RNA for Iso-Seq sequencing. JG cleaned Iso-Seq data, obtained concensus circular sequences (CCS) and assembled the Ha7650b/ PI413066 locus. HB developped the bioinformatic pipeline for isoseq scan. HB and AF performed the isoseq scan and in-depth analysis of candidate sequences. NP designed primers for PCR amplification and Sanger sequencing of candidates sequences and performed the amplifications. NP and HB analysed Sanger sequences. HB funded the study. HB and SM coordinated the study. HB wrote the manuscript.

