## Supplementary Tables 1-8 ; Supplementary Methods for "Pooled Single-Molecule transcriptomics identifies a giant gene under balancing selection in sunflower"

### Supplementary Information

#### Supplementary Tables

**Supplementary Table 1:** Raw data from two baches of Iso-Seq sequencing.

| | Batch 1 | | Batch 2 | |
|---|---|---|---|---|
| SMRT-cell ID | A01@044 | A01@045 | A01@052 | A01@053 |
| Number of Bases | 3.89 Gbp | 6.21 Gbp | 11.66 Gbp | 8.85 Gbp |
| Number of Reads | 136,172 | 202,886 | 735,395 | 598,254 |
| Mean Read Length | 31,653 bp | 31,706 bp | 15,929 bp | 14,904 bp |
| N50 Read Length | 54,750 bp | 56,750 bp | 39,250 bp | 35,250 bp |
| Number of SubReads | 2,216,437 | 3,545,193 | 9,330,346 | 7,114,953 |
| N50 SubRead Length | 2,750 bp | 2,750 bp | 2,250 bp | 2,250 bp |

**Supplementary Table 2:** Summarized statistics of consensus circular sequences (CCS) obtained from subreads. cDNA libraries were constructed from a pool of pistil mRNA sampled from 8 individuals of the same wild population of *Helianthus annuus*. Each batch consists in the sequencing of two SMRT cells. NUM = number of sequences. BP = total number of nucleotides.

| | | |
|---|---|---|
| NUM | 188,154 | 519,734 |
| MIN | 26 | 0 |
| MAX | 15,411 | 15,455 |
| N50 BP | 2,196 | 1,723 |
| N50 NUM | 64,011 | 172,076 |
| N90 BP | 1,368 | 1,012 |
| N90 NUM | 150,415 | 394,497 |
| MEAN | 2,032 | 1,491 |
| MEDIAN | 1,873 | 1,471 |
| BP | 382,422,255 | 775,061,062 |

**Supplementary Table 3:** Characteristics of the 6 main outlier genes for nucleotide and haplotypic diversity (CCS = consensus circular sequences)

| Cluster ID | Chr | Start | Number of CCS | Annotation | Number of divergent haplotypes | π | $\pi_N/\pi_S$ | Overlapping QTL for SI |
|---|---|---|---|---|---|---|---|---|
| 1410 | 2 | 4,309,587 | 80 | Serine/threonine receptor kinase | 5 | 0.0788 | 0.594 | No |
| 7867 | 7 | 9,396,775 | 19 | Fbox leucine-rich repeat | 3 | 0.114 | 0.622 | No |
| 8868 | 8 | 8,010,685 | 15 | Toll-like receptor | 6 | 0.094 | 0.655 | No |
| 19877 | 15 | 2,615,002 | 24 | Ubiquitin | 5 | 0.089 | 0.006 | No |
| 23856 | 17 | 186,562,151 | 20 | Serine/threonine receptor kinase | 4* | 0.106 | 0.490 | Yes |
| 24756 | 4 | 16,609,506 | 9 | Non-coding | 4 | 0.0694 | NA | No |

* one of the four divergent haplotype included two closely-related alleles, S4-S6, see Figure 3.

**Supplementary Table 4**: Re-mapping of the QTL for SI loss against the sunflower reference genome (Chr = chromosome)

| marker | distance to SI QTL | database | Accession_number | Chr | Start | End | identity |
|---|---|---|---|---|---|---|---|
| HT945 | 6.1 cM | sunflower transcriptome database* | Ha412T4l22256C0S1 | chr17 | 176,199,443 | 176,199,076 | 96.2% |
| ORS735 | 2.4 cM | GenBank | BV006734.1 | chr17 | 191,197,521 | 191,197,924 | 98.8% |

* https://www.sunflowergenome.org

**Supplementary Table 5:** List of consensus circular sequences corresponding to alleles and isoforms of *Hea7650b*. SX refers to the allele (X) and iN refers to the isoform (N).

| CSS_ID | Allele_ID |
|---|---|
| 54161_180810_091218/17040027/ccs | S6i1 |
| 54161_180810_091218/68682525/ccs | S6i2 |
| 54161_180808_090256/23396777/ccs | S6i3 |
| 54161_181016_141058/16057164/ccs | S6i4 |
| 54161_181016_141058/12714596/ccs | S6i5 |
| 54161_180808_090256/36176199/ccs | S6i6 |
| 54161_180810_091218/11338207/ccs | S5i1 |
| 54161_180810_091218/44302977/ccs | S5i2 |
| 54161_181018_125601/21954737/ccs | S5i3 |
| 54161_181016_141058/13763156/ccs | S5i4 |
| 54161_181016_141058/43058104/ccs | S5i5 |
| 54161_181016_141058/53608563/cc | S4i1 |
| 54161_181018_125601/42074287/ccs | S4i2 |
| 54161_180810_091218/30671234/ccs | S3i1 |
| 54161_180810_091218/45875328/ccs | S3i2 |
| 54161_181018_125601/63635896/ccs | S3i3 |
| 54161_180810_091218/54591635/ccs | S2i1 |
| 54161_181016_141058/54395686/ccs | S2i2 |
| 54161_181018_125601/8520130/ccs | S1i1 |
| 54161_181016_141058/37814935/ccs | S1i2 |
| 54161_181016_141058/55378221/ccs | S1i3 |
| 54161_180810_091218/6291896/ccs | S1i4 |
| 54161_181018_125601/51708828/ccs | S1i5 |
| 54161_180808_090256/21234302/ccs | S1i6 |
| 54161_181018_125601/49545341/ccs | S1i7 |

**Supplementary Table 6:** Number of segregating sites in the total alignment of the CDS of Hea7650b or in CDS regions corresponding to the LRR, malectin, transmembrane and kinase domains of the gene product. Either all 25 CCS covering the gene were used, or a consensus sequences of each allele (6 alleles).

| | Total alignment | | LRR | | Malectin | | Transmembrane | | Kinase | |
|---|---|---|---|---|---|---|---|---|---|---|
| Nb sites | 25 CCS | 6 alleles | 25 CCS | 6 alleles | 25 CCS | 6 alleles | 25 CCS | 6 alleles | 25 CCS | 6 alleles |
| monomorphic | 2,055 | 2,15 | 657 | 685 | 369 | 393 | 51 | 53 | 718 | 747 |
| bi-allelic | 819 | 751 | 343 | 322 | 150 | 132 | 13 | 11 | 200 | 177 |
| tri-allelic | 139 | 115 | 75 | 70 | 33 | 27 | 3 | 3 | 11 | 5 |
| quadri-allelic | 13 | 10 | 10 | 8 | 2 | 2 | 0 | 0 | 0 | 0 |
| length | 3,026 | 3,026 | 1,085 | 1,085 | 554 | 554 | 67 | 67 | 929 | 929 |

**Supplementary Table 7:** Rates of segregating sites in the total alignment of the CDS of Hea7650b or in CDS regions corresponding to the LRR, malectin, transmembrane and kinase domains of the gene product. Either all 25 CCS covering the gene were used, or a consensus sequences of each allele (6 alleles).

| | Total alignment | | LRR | | Malectin | | Transmembrane | | Kinase | |
|---|---|---|---|---|---|---|---|---|---|---|
| Nb sites | 25 CCS | 6 alleles | 25 CCS | 6 alleles | 25 CCS | 6 alleles | 25 CCS | 6 alleles | 25 CCS | 6 alleles |
| monomorphic | 0,679 | 0,711 | 0,606 | 0,631 | 0,666 | 0,709 | 0,761 | 0,791 | 0,773 | 0,804 |
| bi-allelic | 0,271 | 0,248 | 0,316 | 0,297 | 0,271 | 0,238 | 0,194 | 0,164 | 0,215 | 0,191 |
| tri-allelic | 0,046 | 0,038 | 0,069 | 0,065 | 0,060 | 0,049 | 0,045 | 0,045 | 0,012 | 0,005 |
| quadri-allelic | 0,004 | 0,003 | 0,009 | 0,007 | 0,004 | 0,004 | 0,000 | 0,000 | 0,000 | 0,000 |

**Supplementary Table 8:** List of sequences used for the phylogenetic analysis of the At1g07650 orthogroup in Asterids.

| | | | | |
|---|---|---|---|---|
| *Helianthus annuus* | www.heliagene.org | HanXRQChr01g0028671 | Hea7650a | Ingroup |
| *Helianthus annuus* | Pooled isoseq | S1 | Hea7650bS1 | Ingroup |
| *Helianthus annuus* | Pooled isoseq | S2 | Hea7650bS2 | Ingroup |
| *Helianthus annuus* | Pooled isoseq | S3 | Hea7650bS3 | Ingroup |
| *Helianthus annuus* | Pooled isoseq | S4 | Hea7650bS4 | Ingroup |
| *Helianthus annuus* | Pooled isoseq | S5 | Hea7650bS5 | Ingroup |
| *Helianthus annuus* | Pooled isoseq | S6 | Hea7650bS6 | Ingroup |
| *Helianthus annuus* | www.heliagene.org | HanXRQChr17g0565721 | Hea7650c | Ingroup |
| *Helianthus annuus* | www.heliagene.org | HanXRQChr17g0565731 | Hea7650d | Ingroup |
| *Helianthus annuus* | www.heliagene.org | HanXRQChr17g0565871 | Hea7650e | Ingroup |
| *Ipomoea nil* | OrthoDB | XM_019338971.1 | Ipn7650 | Ingroup |
| *Capsicum annuum* | OrthoDB | PHT76856 | Caa7650 | Ingroup |
| *Capsicum baccatum* | OrthoDB | PHT43633 | Cab7650 | Ingroup |
| *Coffea arabica* | OrthoDB | XM_027205493 | Coa7650 | Ingroup |
| *Camellia sinensis* | OrthoDB | XM_028244167 | Cas7650 | Ingroup |
| *Lactuca sativa* | OrthoDB | XM_023910336.1 | Las7650 | Ingroup |
| *Mikania micrantha* | Mikania micrantha genome | KAD4982895.1 | Mim7650a | Ingroup |
| *Mikania micrantha* | Mikania micrantha genome | Manual annotation | Mim7650b | Ingroup |
| *Nicotiana attenuata* | OrthoDB | XM_019368822.1 | Nia7650 | Ingroup |
| *Artemisia annua* | Artemisia genome | PKPP01000279 | Ara7650 | Ingroup |
| *Cynara cardunculus* | OrthoDB | XM_025131696 | Cyc7650 | Ingroup |
| *Daucus carota* | OrthoDB | XM_017390532 | Dac7650 | Ingroup |
| *Chrysanthemum nankingense* | Chrysantenum genome | CHR00029983-RA | Chr7650 | Ingroup |
| *Olea europaea var. sylvestris* | OrthoDB | XM_022992872.1 | Ole7650 | Ingroup |
| *Sesamum indicum* | OrthoDB | XM_011099761.2 | Sei7650 | Ingroup |
| *Solanum lycopersicum* | OrthoDB | XM_004243193.3 | Sol7650 | Ingroup |
| *Solanum pennellii* | OrthoDB | XM_015227316.2 | Sop7650 | Ingroup |
| *Solanum tuberosum* | OrthoDB | XM_006348867.2 | Sot7650 | Ingroup |
| *Populus trichocarpa* | OrthoDB | XM_024599333.1 | Pot7650 | Outgroup |

### Supplementary Methods

##### *Allelic amplifications by long-range PCR*

PCR amplifications were carried out in 96-well plates with a total volume of 17µl. PCR reaction for full-length cDNA amplification or 1-kb amplifications was 20ng of 1/20 diluted cDNA, 1X HF buffer (Thermo Fisher Scientific, USA), 150µM dNTP (Promega, USA), 0.2µM of forward and reverse primers and 0.2U of Phusion High-Fidelity DNA Polymerase (Thermo Fisher Scientific, USA). Amplifications were performed with a touchdown PCR protocol: after an initial 1min pre-incubation step at 98°C, 11 cycles each of 10s denaturation at 98°C, 30s at the annealing temperature which decreased by 0.6°C per cycle from 64°C to 56°C, and 30s (for 1kb-amplicons) or 2min (for full-length cDNA amplifications) elongation at 72°C. The program was followed by 25 additional cycles, each consisting of 10s at 98°C, and 30s at 56°C and 30s or 2min at 72°C, ended by a 5min elongation step at 72°C. Amplification products were separated on 2% agarose gels buffered with 0.5X TAE and visualized by ultraviolet illumination after ethidium bromide staining. Amplicons purifications for sequencing were performed with 1% Sera-Mag SpeedBeads™ Carboxyl Magnetic Beads (GE Healthcare, USA), with 2 washes with 70% ethanol and elution in 20µl 10/0.1 TE buffer. Purified amplicons has been submitted to Eurofins Genomics, Germany, for Sanger sequencing.

##### *Sequencing and assembly of the Hea7650b locus of the PI413066 wild sunflower from New Mexico*

First the reads were assembled using CANU[54] version 1.9 with stringent parameters (minReadLength=5000, utgErrorRate=0.0100, batOptions=-dg 3 -db 3 -dr 1 -ca 500 -cp 50, genomeSize=3g -pacbio-hifi). The CANU assembly spans 3,905,987,352 bp in 20,351 contigs (N50: 533,277bp). The mapping of mRNAs allowed us to identify that the locus was split in three contigs (tig00002282:1,992,059nt; tig00009922:238,489nt; tig00010878: 251,056nt). We were unable to find CANU parameters able to solve the assembly of the locus in a contiguous sequence. In order to test alternate algorithm, the FALCON[55] pipeline was run on the same dataset (version pb-assembly_201912, pa_daligner_option = -B24 -e.80 -l1000 -s1000 -M42; ovlp_daligner_option = -B24 -h60 -e.96 -l500 -s1000 -M42; ovlp_HPCdaligner_option = -B24 -h60 -e.96 -l500 -s1000 -M42; pa_DBsplit_option = -x500 -s200; ovlp_DBsplit_option = -x500 -s200). The overlap filtering step of the FALCON pipeline was replaced by til-r[56] software using stringent parameters (version 20180523; parameters: --

minwing 500 --minovl 2500 --minpci 98 --deltapci 1). The FALCON assembly spans 2,738,472,176 bp in 5287 contigs (N50: 1,348,425bp). None single falcon contig spans the whole locus. But, as the falcon contig 001915F (392,809nt) overlaps the CANU contig tig00002282, the two contigs were assembled using minimus2 (default parameters, version amos-3.1.0). Minimus2 found an overlap of 60,821nt at 99.82 percent identity and generated a consensus sequence spanning the whole locus in a single contiguous sequence. The mRNA sequence data were mapped using gmap (version 2019-09-12, same parameters as above) to annotate a gene spanning 385,712 nucleotides in the PI413066 wild sunflower.
